# Feeling left out or just surprised? Neural correlates of social exclusion and over-inclusion in adolescence

**DOI:** 10.1101/524934

**Authors:** Theresa W. Cheng, Nandita Vijayakumar, John C. Flournoy, Zdena Op de Macks, Shannon J. Peake, Jessica E. Flannery, Arian Mobasser, Sarah L. Alberti, Philip A. Fisher, Jennifer H. Pfeifer

**Affiliations:** University of Oregon, Eugene, Oregon, USA; Deakin University, Melbourne, Australia; Harvard University, Cambridge, Massachusetts, USA; Leiden University, the Netherlands; Oregon Health & Science University, Portland, Oregon, USA

**Keywords:** Cyberball, social exclusion, functional MRI, anterior cingulate cortex, anterior insula

## Abstract

Social belonging is an important human drive that influences mood and behavior. Neural responses to social exclusion are well-characterized, but the specificity of these responses to processing rejection-related affective distress is unknown. The present study compares neural responses to exclusion and over-inclusion, a condition that similarly violates fairness expectations but does not involve rejection, with a focus on implications for models of dorsal anterior cingulate cortex (dACC) and anterior insula (AI) function. In an fMRI adaptation of the Cyberball paradigm with adolescents 11.1-17.7 years of age (N=69), we employed parametric modulators to examine scaling of neural signal with cumulative exclusion and inclusion events, an approach that overcomes arbitrary definitions of condition onsets/offsets imposed on fluid, continuous gameplay. We identified positive scaling of dACC and posterior insula response with cumulative exclusion events, but these same regions exhibited trending signal *decreases* with cumulative inclusion events. Furthermore, areas within the dACC and insula also responded to context incongruency (throws to the participant in the exclusion run; throws between computer players in the over-inclusion run). Taken together, these findings caution against interpretations that responses in these regions uniquely reflect aspects of affective distress within social rejection paradigms. We further identified that the left ventrolateral PFC, rostromedial PFC, and left intraparietal sulcus responded similarly to cumulative exclusion and inclusion. These findings shed light on which neural regions exhibit patterns of differential sensitivity to exclusion or over-inclusion, as well as those that are more broadly engaged by both types of social interaction.

## 1. Introduction

From the earliest fMRI studies on social rejection, it has been proposed that the dorsal anterior cingulate cortex (dACC) and anterior insula (AI) process the affective distress underlying both physical and social pain (Eisenberger, Lieberman, & Williams, 2003; Eisenberger, Jarcho, Lieberman, & Naliboff, 2006). This hypothesis suggests that neural resources dedicated to pain processing were co-opted over the course of evolution to make the experience of social rejection particularly salient, motivating the maintenance of close social ties and ultimately promoting evolutionary fitness (Baumeister and Leary, 1995; Eisenberger, 2012). While recent studies suggest that pain and social rejection have distinct representations in the dACC and AI (Woo et al., 2014; Kragel et al., 2018), many perspectives maintain a role for these regions in processing rejection-related affective distress.

This hypothesis can be evaluated by considering whether a brain regions’ responses are specific to social rejection when compared to social over-inclusion. Cyberball (Williams, Cheung, & Choi, 2000) is a virtual ball-tossing game commonly used to simulate social exclusion in neuroimaging studies (for review and meta-analysis, see Vijayakumar, Cheng, & Pfeifer, 2017). Such studies often compare social exclusion to a condition referred to as “social inclusion” that resembles fair play (i.e. roughly equal participation by all players). However, this comparison forms a weak basis for claiming that a region is involved in processing affective distress, because exclusion differs from fair play by defying widely-held baseline expectations of approximate fairness (Gunther Moor et al., 2010; Rodman et al., 2017), thereby inducing expectancy violations. Over-inclusion may be a more appropriate comparison for determining the specificity of neural responses to affective distress, because its psychological experience may be better matched in the degree to which participant’s expectations are violated (Kawamoto et al., 2012). Over-inclusion may also be more similar to exclusion than fair play on other dimensions, such as the intensity of the affective experience. However, few human neuroimaging studies compare social exclusion to over-inclusion.

Existing studies that compare social exclusion with over-inclusion and/or acceptance do not paint a consistent picture of dACC and AI involvement. One such study with Cyberball in young adults found that the dACC exhibited greater responses to exclusion than over-inclusion, but that the AI did not (Kawamoto et al., 2012). In contrast, a social judgment study in late adolescents/young adults found that the dACC and left AI responded to both positive and negative social feedback (Dalgleish et al., 2017). Using a similar paradigm, a study in young adults found that the ventral (v)ACC responded to positive social feedback, while the dACC responded to expectancy violations across positive and negative feedback conditions (Somerville et al., 2006).

Moreover, inconsistencies in the role of the ACC and insula are also evident in quantitative meta-analyses examining neural responses to social exclusion across different paradigms. Of the three meta-analyses, only one identified involvement of the dACC during social exclusion (Rotge et al., 2015). In comparison, more ventral regions of ACC were more reliably recruited across the meta-analyses, including (1) rostral perigenual ACC bordering on mPFC (Cacioppo et al., 2013), (2) perigenual and subgenual ACC (Vijayakumar et al., 2017), and (3) vACC (Rotge et al., 2015). Of the two whole-brain meta-analyses, the AI was identified by one (Cacioppo et al., 2013), but not the other (Vijayakumar et al., 2017). In contrast, both of these studies suggest more reliable engagement of the ventromedial PFC and ventrolateral PFC/lateral OFC. Whether the involvement of these other regions reflects processes unique to social rejection or whether they would also be identified during over-inclusion remains unknown.

Therefore, the present study compares social exclusion to over-inclusion. We employ an fMRI adaptation of Cyberball containing periods of over-inclusion and exclusion interspersed with periods of fair play. This strategy was employed by Kawamoto and colleagues (2012), although they were constrained by arbitrary operationalization of inclusion and exclusion on continuous gameplay (e.g., the N^th^ throw between computer players marks the onset of exclusion). We reduce such constraints by using parametric modulators to analyze changes in BOLD signal associated with cumulative exclusion and inclusion events. Given considerable interest in the role of the dACC and evidence from meta-analyses that the broader extent of the ACC is relevant to these processes, we interrogate eight parcels along the ACC as regions-of-interest (ROIs). We examine five additional ROIs reflecting both regions of similar theoretical interest (left and right AI) and regions that have been reliably recruited in Cyberball paradigms in a prior meta-analysis (left inferior frontal gyrus, posterior cingulate cortex, and ventral striatum, abbreviated VS; Vijayakumar et al., 2017).

Patterns of signal in the dACC and AI are evaluated for consistency with potential roles in processing affective distress and expectancy violation. Based on the assumption that exclusion, but not over-inclusion, induces rejection-related affective distress, we interpret affective distress models to suggest that the dACC and AI will exhibit greater signal scaling with cumulative exclusion events than inclusion events, and that this difference will be driven by signal increases to cumulative exclusion only. Based on the assumption that both exclusion and over-inclusion similarly violate fairness expectations, we interpret expectancy violation accounts to suggest that signal will scale similarly across both conditions. We take another modeling approach to explore neural responses to short-term expectancy violations, i.e. when a ball toss including or excluding the participant is incongruent with the broader inclusionary or exclusionary context established by the run.

Finally, we also examine age-related differences in our sample of 11-17 year-olds. Developing the capacity to navigate complex social relationships is a critical aspect of adolescence (Blakemore & Mills, 2014). Social evaluation becomes a dominant concern (Somerville, 2013), with adolescents spending a significant amount of time with peers both on and offline (Csikszentmihalyi & Larson, 1984; Anderson & Jiang, 2018). Accordingly, adolescents experience stronger affective responses to negative social feedback than adults (Sebastian, Viding, Williams, & Blakemore, 2010). The VS plays a role in affective reactivity (Knutson, Westdorp, Kaiser, & Hommer, 2000; Sescousse, Caldu, Segura, & Dreher, 2013; Jensen et al., 2003; Levita et al., 2009), and its responses to social stimuli are negatively associated with peer susceptibility in adolescence (Pfeifer et al., 2011). Based on a prior meta-analysis identifying this region as more reliably recruited in Cyberball for adolescents than adults (Vijayakumar et al., 2017), we hypothesize that we will identify an age-related decrease in VS response.

## 2. Materials & Methods

### 2.1 Participants

Ninety-seven adolescents ages 11 to 17 were recruited from Lane County, Oregon. These participants form the community control group in a study that included additional samples from the child welfare and juvenile justice systems. Anticipating that mental health diagnoses might be prevalent in these additional samples, we did not exclude for reported current or prior diagnoses of common psychiatric disorders in order to improve comparability across samples. These disorders included anxiety, depression, attention deficit disorder/attention deficit hyperactivity disorder, oppositional defiant disorder, or conduct disorder. However, no participants from the child welfare and juvenile justice system samples are included in any analyses presented in this paper. Of the 97 participants recruited from the community sample only, nine dropped out of the study, seven elected not to participate in the MRI portion, eight completed an alternate (pilot) version of the Cyberball task, one failed to complete the Cyberball task, and one participant’s data was not collected due to technical errors. Additionally, two participants were excluded when visual quality inspection of the imaging data revealed extreme motion and/or orbitofrontal signal dropout.

The analyses presented here were conducted with the remaining 69 participants (36 female) from the community sample aged 11 to 17 years (range= 11.1 to 17.7, M=14.2, SD=1.5). The sample size was based on a priori power calculations for a different research question and statistical approach (group comparisons with the child welfare and juvenile justice system samples). Post-hoc analyses suggest that we have sufficient power (> 0.8) to detect effect sizes of 0.31 or greater in two-tailed one-sample t-tests (e.g. Increasing Exclusion = 0) and effect sizes of .35 or greater in one-tailed one-sample t-tests of the difference between two dependent means (e.g. Increasing Exclusion > Increasing Inclusion).

Our final sample included four subjects that had disclosed psychiatric diagnoses, including two with anxiety disorders and two with attention-related disorders. Six participants reported taking medication for behavioral issues (four subjects reported taking anti-depressant medications, one reported taking a mood stabilizer, and one reported taking stimulants typically prescribed for attention disorders). In total, eight participants reported either receiving psychiatric diagnoses or taking psychotropic medications. We conducted sensitivity analyses by re-running ROI and whole-brain analyses with the parametric modulators in a sample that excluded these eight participants (N = 61).

### 2.2 Procedures

#### 2.2.1 Study Procedure

Written informed consent was obtained from the parent/guardian, while written assent was obtained from the adolescent. At the first visit, the adolescent was screened for MRI eligibility as per procedures determined by the University of Oregon’s Lewis Center for Neuroimaging. The second visit was typically scheduled within one month of the first visit (M=14.8 days, SD=16.5, range=1 to 106). Eligible participants then completed the MRI portion of the study as well as additional questionnaires and behavioral tasks, including the vocabulary and matrix reasoning subtests of the Wechsler Abbreviated Scale of Intelligence, second edition (Wechsler, 1999). Participants were compensated for their time, and all materials and procedures were approved by the Institutional Review Board at the University of Oregon.

#### 2.2.2 The Cyberball task

Developed by Williams, Cheung, and Choi (2000), Cyberball is an animated interactive ball-tossing computer game used to virtually simulate the experience of social exclusion. In our study, participants were informed that they were playing Cyberball with two adolescent peers at partner sites in real time via the Internet. However, the other player throws were computer-automated. Similar adaptations of Cyberball have successfully simulated peer rejection in adolescents by leading them to believe that computer players were real peers (Bolling et al., 2011a; Masten et al., 2009). Participants also made short video profiles to introduce themselves to the other players and viewed the other players’ profiles, further lending credibility to the cover story; to our knowledge, this has not been done in prior studies. Participants were instructed on how to play Cyberball in a mock scan session immediately prior to their MRI session.

Participants played an over-inclusion and exclusion run of Cyberball. Each run contained a total of 44 ball throws. They were instructed to use a button box to indicate which of the two players they wanted to throw to, and if they did not make a decision within 5 s, the ball would be automatically thrown to another player at random. Traditional Cyberball paradigms seek to induce strong feelings of rejection and thus consist of extended periods of exclusion, sometimes taking up close to the entirety of the run. We instead employed periods of exclusion or over-inclusion interspersed with periods of fair play to reflect more fluid interactions. In the inclusion run, participants experienced periods of over-inclusion in which computer players repeatedly threw the ball to the participant rather than to one another. Periods of over-inclusion were interspersed with periods of fair play such that, overall, 17 of the 27 throws by the other players were to the participant (63%). In the exclusion run, participants experienced periods in which the computer players only threw the ball to one another and not to the participant. These were interspersed with periods of fair play such that, overall, six of the other players’ total combined 38 throws were to the participant (16%). In social judgment and evaluation tasks, adolescents’ (ages 11-17) mean expectations of being liked are between 40-55%, and on average, such ratings are lower in mid-adolescence, i.e., between the ages of 12-14 (Gunther Moor et al., 2010; Rodman et al., 2017). Although our choice of paradigm differs, we suggest that the proportion of throws to the participant (63% in over-inclusion and 16% in exclusion) was extreme enough to elicit the intended social manipulation in each case. (Note that these percentages are true of the overall run, and within each run participants experienced even more extreme periods of over-inclusion or exclusion interspersed with periods of fair play). As other computer players still both received the ball some of the time and at approximately even rates, it is generally the case that neither computer player was explicitly excluded by greater participant inclusion. The time elapsed between each computer player catching and throwing the ball was a random number between 0 and 3 s (M=1.5, SD=0.9), and the ball took approximately 1.4 s to travel through the air. Therefore, the timing of events, including participant button presses, varied from trial to trial and did not systematically align with the TRs. See Figure 1 for a schedule of throws in both runs.

**Figure 1.**
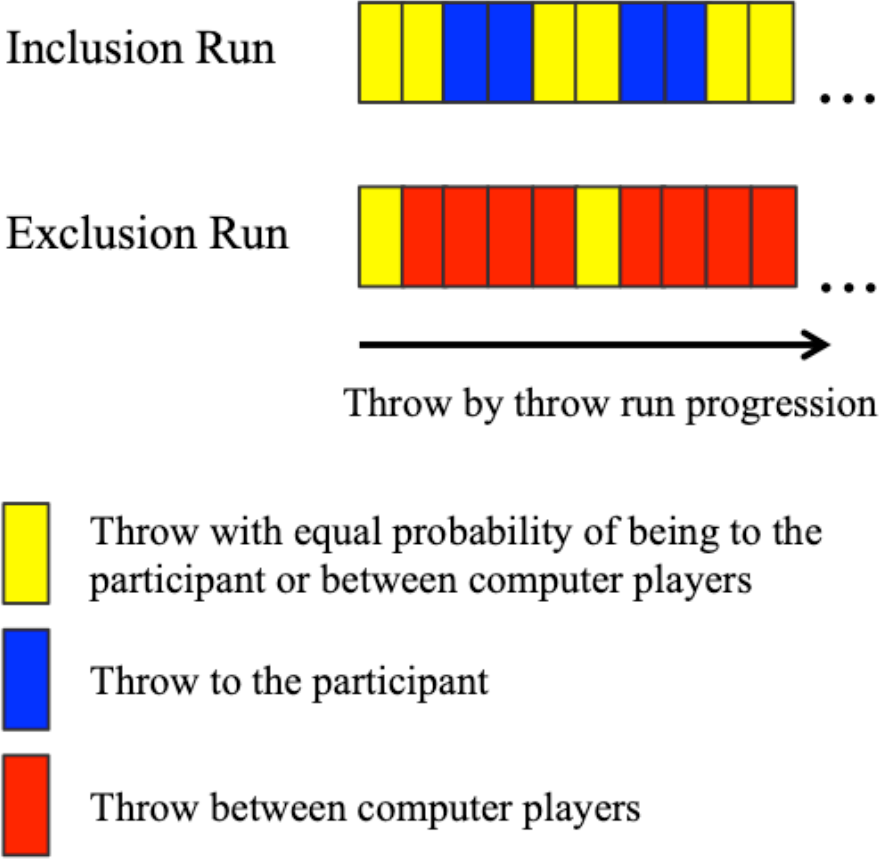
Partial schedules of throws for each of the computer players in both runs are displayed. Each rectangle represents one throw by the computer player. The color of the rectangle indicates the recipient of the ball. The complete schedules extend the pattern displayed to the total number of computer-programmed throws per run, and both of the computer players follow the same schedule within a run.

In the protocol for this study, participants first played two rounds of the Yellow Light Game (YLG), a driving simulation adapted from the Stoplight Task (Gardner & Steinberg, 2005) outside of the scanner. Afterward, within the scanner, participants (a) played two rounds of the YLG, (b) played the Cyberball inclusion run, (c) were introduced to virtual peers and observed them playing the YLG, (d) played two peer-observed runs of YLG, (e) played the Cyberball exclusion run, and (f) played two final peer-observed runs of YLG. The total length of the scan protocol was 1 hr 15 min. The inclusion run always preceded the exclusion run; Cyberball conditions were not counterbalanced due to concern about carryover of negative affect following exclusion (see section 4.5 for a discussion of order effects). Participants were led to believe that peers from the YLG observations were the same as those in the Cyberball game. More details about this task and the overall protocol can be found in work by Op de Macks and colleagues (2018).

#### 2.2.3 Behavioral measures

Following the MRI scan (and typically within 30 minutes of the end of the scan), adolescents completed the Need-Threat Scale (NTS) to assess their subjective experience of distress during Cyberball. We used the 12-item version (Zadro et al., 2004), which has been validated as an overall measure of need-threat (Gerber et al., 2017). The 12 items on the NTS included ratings of belongingness, control, meaningfulness, and self-esteem on a scale from 1 (“not at all”) to 5 (“very much so”), with higher scores reflecting more threat. The overall scale exhibited good reliability in our sample (standardized Cronbach’s α=0.81). We assessed the believability of the deception by asking participants “Did you think the peers could actually see you playing [the driving game]?”. This question was part of a task experience survey administered at the very end of the session, just prior to participant debriefing. From this self-report, we inferred whether or not participants believed they were interacting with real peers during Cyberball.

#### 2.2.4 fMRI data acquisition

Data were acquired on a 3T Siemens Skyra MRI scanner at the Lewis Center for Neuroimaging at the University of Oregon. High-resolution T1-weighted structural images were collected with the MP-RAGE sequence (TE=3.41 ms, TR=2500 ms, flip angle=7**°**, 1.0 mm slice thickness, matrix size=256 x 256, FOV=256 mm, 176 slices, bandwidth=190 Hz/pixel). Two functional runs of T2*-weighted BOLD-EPI images were acquired with a gradient echo sequence (TE=27 ms, TR=2000 ms, flip angle = 90**°**, 2.0 mm slice thickness, matrix size=100 x 100, FOV=200mm, 72 slices, bandwidth=1786 Hz/pixel). There were 60 to 87 images per run, as run length varied with participants’ response times during Cyberball. To correct for local magnetic field inhomogeneities, a field map was also collected (TE=4.37 ms, TR=639.0 ms, flip angle=60°, 2.0 mm slice thickness, matrix size=100 x100, FOV=200 mm, 72 slices, bandwidth=1515 Hz/pixel).

#### 2.2.5 fMRI processing

Raw DICOM image files were converted to the NifTI format with MRIConvert (Smith, 2011). Data were preprocessed using Statistical Parametric Mapping software (SPM12, Wellcome Department of Cognitive Neurology, London, UK). Participants’ anatomical images were coregistered to the 152 Montreal Neurological Institute stereotaxic template, segmented into six tissue types, and used to create a group anatomical template using DARTEL. Next, functional images were unwarped using field maps and corrected for head motion via image realignment. A group averaged field map was created and used as a substitute for two participants: one that did not have a field map and another whose fieldmap was not well-aligned with their functional volumes. Unwarped and realigned mean functional images were coregistered to the anatomical image (that had in turn been coregistered to the MNI template). Transformations were applied to warp the functional data to the DARTEL group template, and these normalized images were smoothed using a 6-mm FWHM Gaussian kernel. Preprocessing scripts used for this analysis are available on GitHub at https://github.com/dsnlab/TDS_scripts/tree/cheng_cyb_main/fMRI/ppc/spm/tds2 (SPM scripts) and https://github.com/dsnlab/TDS_scripts/tree/cheng_cyb_main/fMRI/ppc/shell/schedule_spm_jobs/tds2 (shell scripts).

Motion artifacts were identified using an in-house automated script that evaluates changes in image intensity relative to the mean across all subjects, as well as volume-to-volume changes in Euclidean distance. This script for this is publically available (Cosme et al., 2018). We refer interested readers to the most recent version (https://github.com/dsnlab/auto-motion), as well as the branch used in this analysis (https://github.com/dsnlab/TDS_scripts/tree/cheng_cyb_main/fMRI/fx/motion/auto-motion). A regressor of no interest was constructed by marking volumes of the following types: (a) volumes with greater than 0.3 mm of motion in Euclidian distance relative to the previous volume, (b) volumes for which the *mean intensity across voxels* was extreme (3 SDs above or 1.5 SDs below) relative to the mean intensity across subjects and runs, and (c) volumes for which the *standard deviation across voxels* was extreme (3 SDs above or below) relative to the mean standard deviation across subjects and runs. The mean intensity and standard deviation scores were calculated across all runs for all subjects, including volumes collected during the YLG and while participants observed others playing the YLG. Volumes immediately preceding and following marked volumes were also flagged. This script identified head motion in 36 out of 138 total Cyberball runs across 69 participants. Of the volumes marked for motion, the script flagged an average of 3.94 (5.7%) volumes per run (SD=3.58, range=1 to 17, or up to 23.3%). Additionally, our models included four motion parameters (absolute distance, absolute rotation, and the first derivatives of each of these values) as regressors of no interest. As mentioned previously, two participants were excluded on the basis of head motion/signal dropout. The first was identified for exclusion based on visual inspection of pre-processed images that revealed extreme signal dropout in the orbitofrontal gyrus. The second participant was identified for exclusion based on initial visual inspection of contrasts from their single-subject models, which revealed a clear and severe pattern of motion-related striping that indicated that their data was contaminated. No participants were excluded for exceeding an *a priori* threshold of >25% of marked motion volumes within a single run.

### 2.3 Experimental Design and Statistical Analysis

Cyberball was modeled as an event-related design. For each participant’s fixed-effects analysis, a general linear model was created with two regressors of interest modeled as zero-duration events: the computer-generated throws were each modeled as either an Inclusion Event (IncEvent, i.e. throws to the participant) or an Exclusion Event (ExcEvent, i.e. throws to the computer players). These events occurred within an Inclusion Context (IncContext, i.e. inclusion run) or an Exclusion Context (ExcContext, i.e. exclusion run). An additional zero-duration event regressor of no interest marked when participants’ computer avatar “caught” the ball, signaling the participant’s turn to throw the ball. This regressor was included to control for neural responses related to participants’ decision-making and subsequent button-press, as has been used in previous studies with event related designs (Bolling et al., 2011b). There was also a “wait” period at the start of each run, consisting of 6 s where the screen displayed the words “Connecting to other players…” along with a progress bar, plus additional time until the first throw of the game (on average 8 s in total).

Parametric modulators were created to track consecutive IncEvents within the IncContext (referred to as Increasing Inclusion) and consecutive ExcEvents within the ExcContext (referred to as Increasing Exclusion). Parametric modulators were not created for ExcEvents in the IncContext or vice versa due to an insufficient number of such events in respective runs. Each parametric modulator was mean-centered relative to the average number of continuous throws of that type (for Increasing Inclusion: *M*=2.73, *SD*=1.82; for Increasing Exclusion: *M*=5.97, *SD*=3.89). The model was convolved with the canonical hemodynamic response function, and parameter estimates from the GLM were used to create six linear contrast images: one for each of the four types of events (*i.* IncEvent in IncContext, *ii.* ExcEvent in IncContext*, iii.* IncEvent in ExcContext, and *iv.* ExcEvent in ExcContext) relative to wait periods (collapsed across both runs), and one for each of two parametric modulators (Increasing Inclusion and Increasing Exclusion).

#### 2.3.1 ROI analyses

##### 2.3.1.1 ROI selection and definition

Due to considerable interest in these regions throughout the literature, we chose to examine the ACC and AI as ROIs. To define ACC ROIs, we used a parcellation scheme (Craddock et al., 2012) derived from cluster analyses of resting state functional neuroimaging scans. We selected eight ACC parcels (six regions, two were represented with separate parcels in each hemisphere) from the 250-parcel brain map (see Figure 2). We also examined left and right AI ROIs that were created from visual inspection of anatomical landmarks (also used in Morelli & Lieberman, 2013; Falk et al., 2014). We were also interested in comparing signal between social exclusion and over-inclusion in regions commonly associated with social exclusion in Cyberball. Therefore, we examined two ROIs identified from a prior meta-analysis as being reliably recruited across Cyberball studies (Vijayakumar et al., 2017). We extracted signal from the full clusters in the left inferior frontal gyrus extending into lateral orbitofrontal cortex (peak MNI coordinate: −46, 32, −10; volume = 1992 mm^3^) and posterior cingulate cortex (peak MNI coordinate: −8, −56, 12; volume = 1344 mm^3^). We also examined the cluster in the VS (peak MNI coordinate: 8, 6, 2; volume = 864 mm^3^), which was more reliably recruited in developmental as compared to emerging adult samples during Cyberball. We did not include ROIs that were not also reliably recruited across Cyberball studies in non-clinical samples, and we did not include one large cluster that overlapped significantly with ACC ROIs from the Craddock parcellation. (See Vijayakumar et al., 2017 for details on how these ROIs were obtained.)

**Figure 2.**
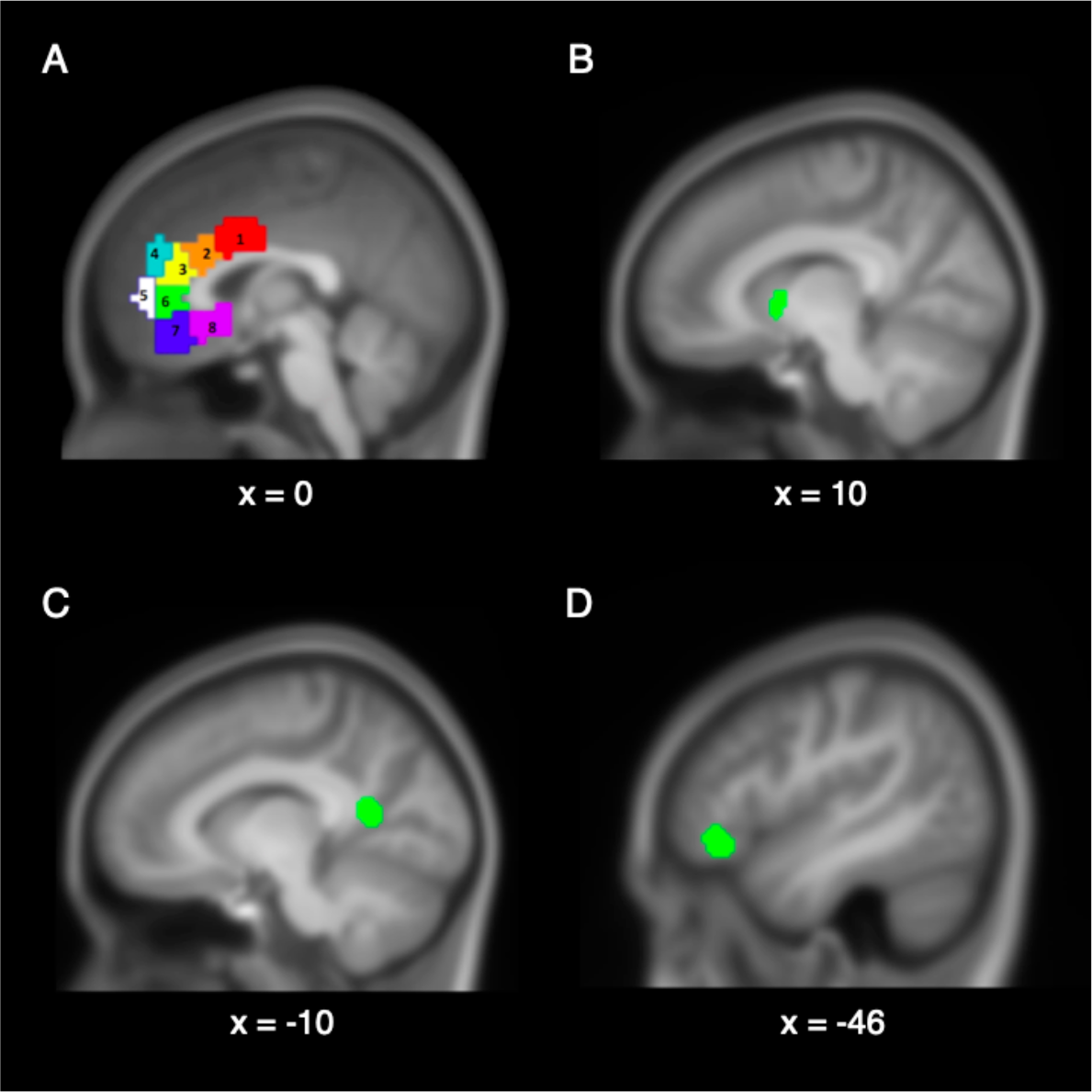
Regions-of-interest (ROIs). (A) Eight ROIs along the extent of the ACC and were identified using the Craddock (2012) parcellation scheme with 250 parcels. (B) The left VS cluster ROI was identified from a meta analysis by Vijayakumar and colleagues (2017) for being more reliably recruited in developmental than emerging adult samples across Cyberball studies. (C) The posterior cingulate cortex cluster and (D) left inferior frontal gyrus cluster ROIs were identified from the same meta-analysis for being reliably recruited across Cyberball studies. The left and right AI ROIs are not pictured here, but can be viewed in Figure 3 of work by Falk and colleagues (2014).

##### 2.3.1.2 ROI extraction and statistical tests

We used MarsBar (version 0.21) to extract parameter estimates of average signal associated with each of the parametric modulators within these parcels. For each parcel, we used a series of t-tests to examine whether signal scaled significantly with cumulative exclusion or inclusion events (null hypotheses: Increasing Inclusion = 0 and Increasing Exclusion = 0). We also examined whether signal scaled differently between cumulative exclusion and inclusion events (null hypothesis: Increasing Inclusion = Increasing Exclusion) via paired, two-tailed student’s t-tests. We controlled the false discovery rate at. 05 using the Benjamini-Hochberg procedure (Benjamini & Hochberg, 1995). We do not apply Bonferroni correction because the set of tests are not independent; adjacent parcels are spatially correlated both because parcels do not necessarily reflect strict signal boundaries and because of smoothing, and the results of different tests are correlated (i.e. if the difference between cumulative exclusion and cumulative inclusion is significantly different, it follows that at least one them is significantly different from zero).

##### 2.3.1.3 Age-related changes in the VS ROI

We used linear regression to examine the association between age and responses to cumulative events in the VS. We regressed parameter estimates averaged across the VS on age for each of three conditions: Increasing Inclusion, Increasing Exclusion, and their difference. For this family of three tests, we accounted for multiple comparisons by using a Bonferroni-corrected threshold of *p*<.0167.

#### 2.3.2 Whole-brain analyses

Using whole-brain analyses, we examined whether smaller clusters within our ROIs reflected important differences between social exclusion and over-inclusion. These clusters may not be identified from ROI analyses due to signal dilution when averaging over larger regions and/or non-optimal parcel boundaries. Additionally, whole-brain analyses were used to identify regions outside of our ROIs that are significantly associated with conditions of interest, particularly for those that are less studied (e.g. Increasing Inclusion). Therefore, we conducted group-level analyses based on these fixed-effects (single subject) contrast images, modeling the subject variable as a random effect. We ran whole-brain conjunction and subtraction analyses with the parametric modulators using paired samples t-tests. We also used a repeated-measures flexible factorial ANOVA with a 2×2 design to examine the interaction between the event (IncEvent and ExcEvent) and context (IncContext and ExcContext) factors. This enabled comparisons between Context Congruent (i.e. IncEvent in IncContext and ExcEvent in ExcContext) and Context Incongruent (i.e. IncEvent in ExcContext and ExcEvent in IncContext) events. The main effects of event and context are also reported in the Supplementary Materials (see section S3). Whole-brain age effects are also reported in the Supplementary Materials (see section S4), and controlling for age did not alter any of the results presented in Tables 3 and 4.

Unless otherwise specified, reported results exceeded the minimum cluster size threshold needed for a .05 family-wise error (FWE) rate given a voxel-wise threshold of p=.001 (bi-sided, NN=3, k=68). Cluster extent thresholds were identified using AFNI 3dClustSim, version AFNI_17.1.01 (Apr 12 2017; version update accounts for software bugs identified by Eklund and colleagues (2016)). Smoothness estimates entered into 3dClustSim were spatial autocorrelation function (acf) parameters averaged from each individual’s first level model residuals as calculated by 3dFWHMx (acf parameters: 0.70986 4.667 8.5925).

#### 2.3.3 Exploratory examination of functionally defined dACC

We used NTS scores to predict the degree of signal scaling to Increasing Exclusion within a functionally-defined dACC ROI. This ROI is a sphere with a radius of 4 mm constructed around the most ventral sub-peak of a dACC-SMA cluster (MNI coordinates: −8, 6, 38) identified from the contrast of Increasing Exclusion > Increasing Inclusion (see Figure 3A). Among cluster sub-peaks, the most ventral peak was visually identified as most clearly within the dACC and thus least likely to include SMA signal. We were interested in whether responses in this region scaled with the affective distress of *exclusion*, and therefore did not examine responses to Increasing Inclusion or the difference between the two conditions. Since only one statistical test was needed to answer this question, we do not correct for multiple comparisons. We additionally visualized the pattern of signal changes to Increasing Exclusion and Increasing Inclusion in this ROI (see Figure 3B).

**Figure 3.**
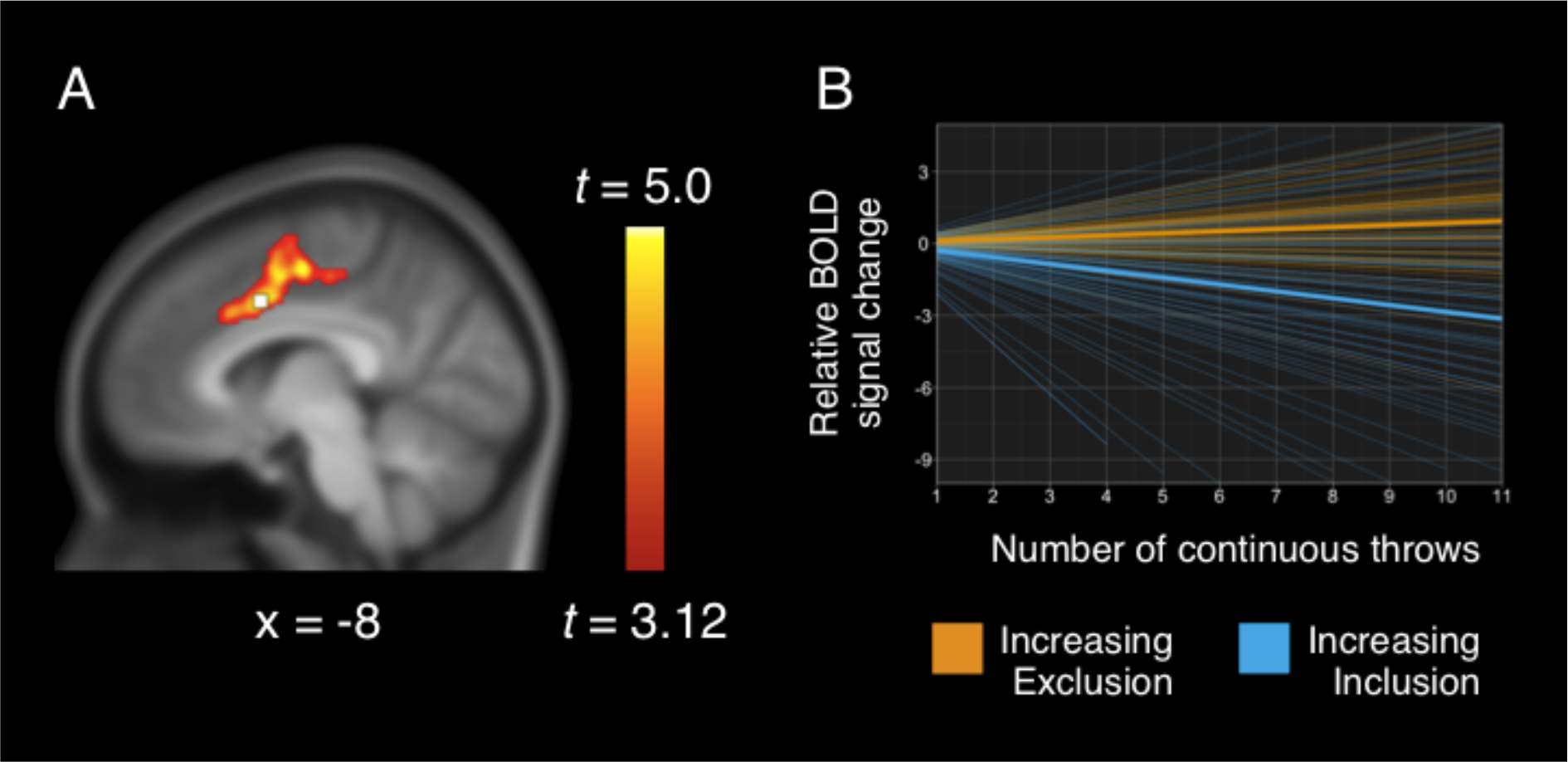
Examination of a functionally-defined dACC ROI. (A) The center of the 4-mm radius sphere was the most ventral sub-peak of a cluster encompassing the dACC identified from the contrast of Increasing Exclusion > Increasing Inclusion. (B) Parameters extracted from this sphere reflect expected signal change per continuous throw, and lines with those slopes are plotted. Mean beta parameter estimates suggest that signal differences between the parametric modulators were driven by decreases with Increasing Inclusion (in yellow, M=-.28, SE=.07), as well as increases with Increasing Exclusion (in blue, M=0.08, SE=.02). Faint lines represent individual estimates, while bold lines reflect the group average.

## 3. Results

### 3.1 Behavioral results

To infer whether or not participants believed they were interacting with real peers during the paradigm, participants were asked in a post-task survey “Did you think the peers could actually see you playing [the driving game]?”. Three participants (4%) did not respond, eight (12%) expressed disbelief, and the remaining majority (58 participants; 84%) believed that they were interacting with real peers. We did not exclude participants based on responses to this survey item because feelings of exclusion can be induced even when participants know they are playing with computer players (Zadro et al., 2004).

On average, participants reported a moderate level of subjective need threat during the Cyberball game (mean NTS score=3.23, SD=.73; higher scores reflect more threat) as measured on the NTS, comparable to levels reported in other studies using Cyberball with adolescents (Masten et al., 2009; Bolling et al., 2011b). One subject did not complete the NTS, and was excluded from subsequent analyses with this scale.

### 3.2 fMRI results

#### 3.2.1. Responses to Increasing Exclusion and Increasing Inclusion within ROIs

See Table 1 for the results of statistical comparisons across ROIs in the ACC, and see Table 2 for results from all other ROIs. Regions-of-interest across the extent of the ACC (ROIs 3, 5, 7, 8), VS, left inferior frontal gyrus and left posterior cingulate cortex exhibited statistically significant increases with Increasing Exclusion after correction for multiple comparisons. Increasing Exclusion was associated with greater signal than Increasing Inclusion in the most superior and caudal dACC parcel only (ROI 1 in Figure 2; *t*(68)=2.97, *p*=.004). This dACC parcel appeared to exhibit both signal increases with Increasing Exclusion (*M*=0.06, *SE*= 0.02, *t(68)*=3.02, *p*=.003) and decreases with Increasing Inclusion (*M*=-0.11, *SE*= 0.05, *t(68)*=-2.06, *p*=.043). However, none of the findings in this ROI were statistically significant after correction for multiple comparisons. The left inferior frontal gyrus also exhibited notable signal increases with Increasing Inclusion that were non-significant after correction for multiple comparisons.

**Table 1.** Evaluating parametric modulators within the ACC regions-of-interest

| ROI | Increasing<br>Exclusion M<br>(SE) | H<sub>0</sub>: Increasing<br>Exclusion = 0<br>*t* (*p*) | Increasing<br>Inclusion M<br>(SE) | H<sub>0</sub>: Increasing<br>Inclusion = 0<br>*t* (*p*) | H<sub>0</sub>: Increasing<br>Exclusion = Increasing<br>Inclusion<br>*t* (*p*) |
| --- | --- | --- | --- | --- | --- |
| 1 | .06 (.02) | 3.02 (3.51e-3)** | -.11 (.05) | -2.06 (.043) | 2.97 (4.04e-3)** |
| 2 | .05 (.02) | 3.09 (2.94e-3)** | -.03 (.06) | -.53 (.598) | 1.40 (.167) |
| 3 | .08 (.02) | 3.69 (4.46e-4)*** | .01 (.08) | 0.12 (.906) | .85 (.396) |
| 4 | .03 (.02) | 1.77 (.081) | .07 (.07) | 1.05 (.296) | -.56 (.579) |
| 5 | .11 (.02) | 4.95 (5.17e-6)*** | .13 (.08) | 1.66 (.101) | -.26 (.792) |
| 6 | .06 (.02) | 3.27 (1.70e-3)** | -.01 (.06) | -.15 (.878) | 1.16 (.249) |
| 7 | .07 (.03) | 2.89 (5.11e-3)** | .11 (.07) | 1.42 (.160) | -.44 (.664) |
| 8 | .12 (.03) | 4.86 (7.12e-6)*** | .22 (.10) | 2.19 (.032) | -.93 (.358) |

*Caption:* M=mean beta, SE= standard error; $H_0$ = null hypothesis; \*\*raw $p < .01$ , \*\*\*raw $p < .001$ , ∙: significant at FDR-adjusted $q = 0.05$ .

**Table 2.** Evaluating parametric modulators within non-ACC regions-of-interest

| ROI | Increasing<br>Exclusion M<br>(SE) | $H_0$: Increasing<br>Exclusion = 0<br>$t(p)$ | Increasing<br>Inclusion M<br>(SE) | $H_0$: Increasing<br>Inclusion = 0<br>$t(p)$ | $H_0$: Increasing<br>Exclusion =<br>Increasing<br>Inclusion<br>$t(p)$ |
| --- | --- | --- | --- | --- | --- |
| Left inferior<br>frontal gyrus | .09 (.02) | 4.91<br>(6.07e-6)*** | .18 (.07) | 2.71 (8.53e-3)** | -1.35 (.181) |
| Left posterior<br>cingulate cortex | .13 (.03) | 4.93<br>(5.55e-6)*** | .02 (.09) | .20 (.839) | 1.14 (.259) |
| VS | .08 (.03) | 3.20 (2.11e-3)** | .24 (.11) | 2.27 (.026) | -1.50 (.139) |
| Left AI | .06 (.02) | 2.95 (4.34e-3)** | .02 (.05) | .35 (.730) | .72 (.474) |
| Right AI | .05 (.02) | 2.69 (8.95e-3)** | .01 (.04) | .25 (.801) | .97 (.336) |

*Caption:* M=Mean beta, SE= Standard error; $H_0$ = null hypothesis; \*\* $p < .01$ , \*\*\* $p < .001$ , ∙: significant at FDR-adjusted $q = 0.05$ .

We conducted sensitivity analyses in two subsamples: 1) removing eight subjects taking that self-reported psychiatric diagnoses and/or taking psychotropic medications (N = 61), and 2) removing eight subjects that indicated disbelief in the manipulation (N=61). The overall pattern of results in the caudal dACC parcel (ROI 1) in each of the subsamples was consistent with the pattern in the full sample (greater signal in Increasing Exclusion than Increasing Inclusion reflecting signal increases with Increasing Exclusion *and* decreases with Increasing Inclusion), although only signal increases with Increasing Exclusion were significant after correction for multiple comparisons. We note that *every* ROI except the most rostro-superior and rostro-inferior ACC parcels (ROIs 4 and 7) exhibited statistically significant increases with Increasing Exclusion across both subsamples. No ROIs exhibited significant differences between Increasing Exclusion and Increasing Inclusion across both subsamples, and this was generally due to either no responses or modest signal increases with Increasing Inclusion (and no notable decreases with Increasing Inclusion except for in the caudal dACC parcel). After excluding those with mental health disorders and/or medications, we also identified significant signal increases to Increasing Inclusion in the VS. Differences in the pattern of ROI results and whole-brain analyses are generally minor, and are reported in the Supplementary Materials (see section S1).

##### 3.2.1.1 Exploratory examination of a functionally-defined dACC ROI

Averaged signal from a dACC ROI (functionally-defined for exhibiting greater signal with Increasing Exclusion than Increasing Inclusion) was positively associated with participant reports of need-threat (B=.06, SE=.03, *t*(66)=1.83, *p*=.072). An examination of Cook’s d statistic suggested the presence of three outliers (N=68, Cook’s d greater than 4/N, or .059). Re-running model statistics without outliers strengthened the effect (*t*(63)=2.51, *p*=.0146). See Figure 3 for a visualization of this ROI and its parameter estimates.

##### 3.2.1.2 Age associations in the VS ROI

While signal in the VS exhibited significant increases with Increasing Exclusion overall, age was associated with reductions in the VS response to Increasing Exclusion only (B=-.03, SE=.017, *t*(67)=-1.99, *p*=.050; not significant after correction for multiple comparisons). An examination of Cook’s d statistic suggested the presence of four outliers (N=69, Cook’s d greater than 4/N, or .058). Re-running model statistics without outliers weakened the effect (*t*(63)= −1.80, *p*=.077).

#### 3.2.2 Whole-brain responses to Increasing Inclusion and Increasing Exclusion

See Table 3 for a summary of whole-brain results involving parametric modulators. Statistical maps are also available on NeuroVault (https://neurovault.org/collections/3794). The conjunction of Increasing Inclusion and Increasing Exclusion identified shared signal decreases in the left intraparietal sulcus, as well as shared increases in the rostromedial PFC, as illustrated in Figure 4. The conjunction of signal positively associated with Increasing Exclusion and negatively associated with Increasing Inclusion identified clusters in the left primary motor cortex, supplementary motor area, and left posterior insula. The contrast of Increasing Inclusion > Increasing Exclusion identified significant clusters in the supplementary motor area, right cuneus, and in a sub-gyral region ventral to the cingulate gyrus. The contrast of Increasing Exclusion > Increasing Inclusion identified the left motor cortex (encompassing pre- and postcentral gyri and extending into the insula), right posterior insula, and supplementary motor area extending into the dACC. There were no age effects, nor were there age interactions with the parametric modulators. Controlling for age did not alter whole-brain findings with the parametric modulators.

**Figure 4.**
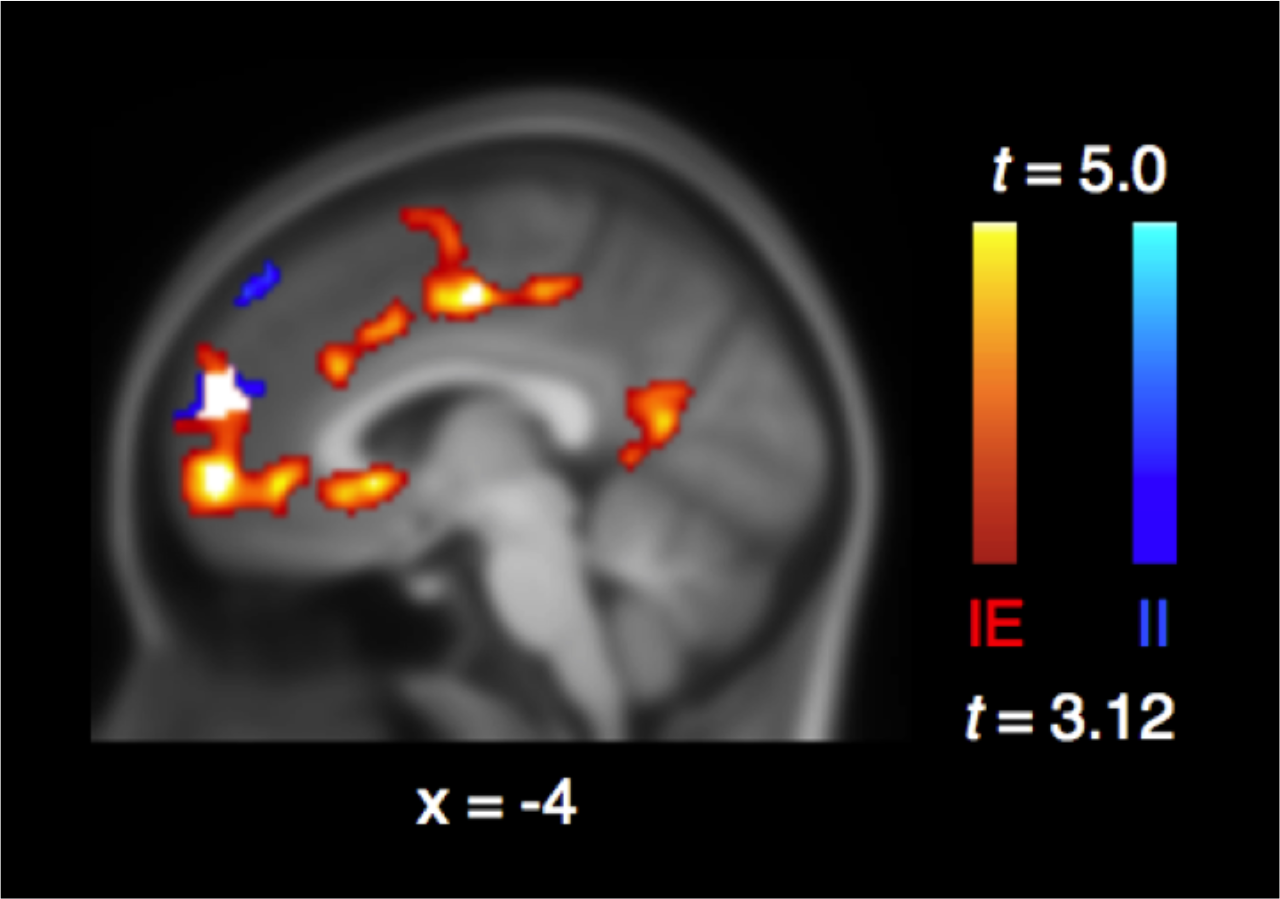
Conjunction of Increasing Exclusion and Increasing Inclusion. The rostromedial prefrontal cortex is identified from the conjunction (white) of parametric modulators for Increasing Exclusion (red) and Increasing Inclusion (blue). Results are FWE cluster-corrected at p<.05 (voxel-wise p<.001, k=68).

**Table 3.**
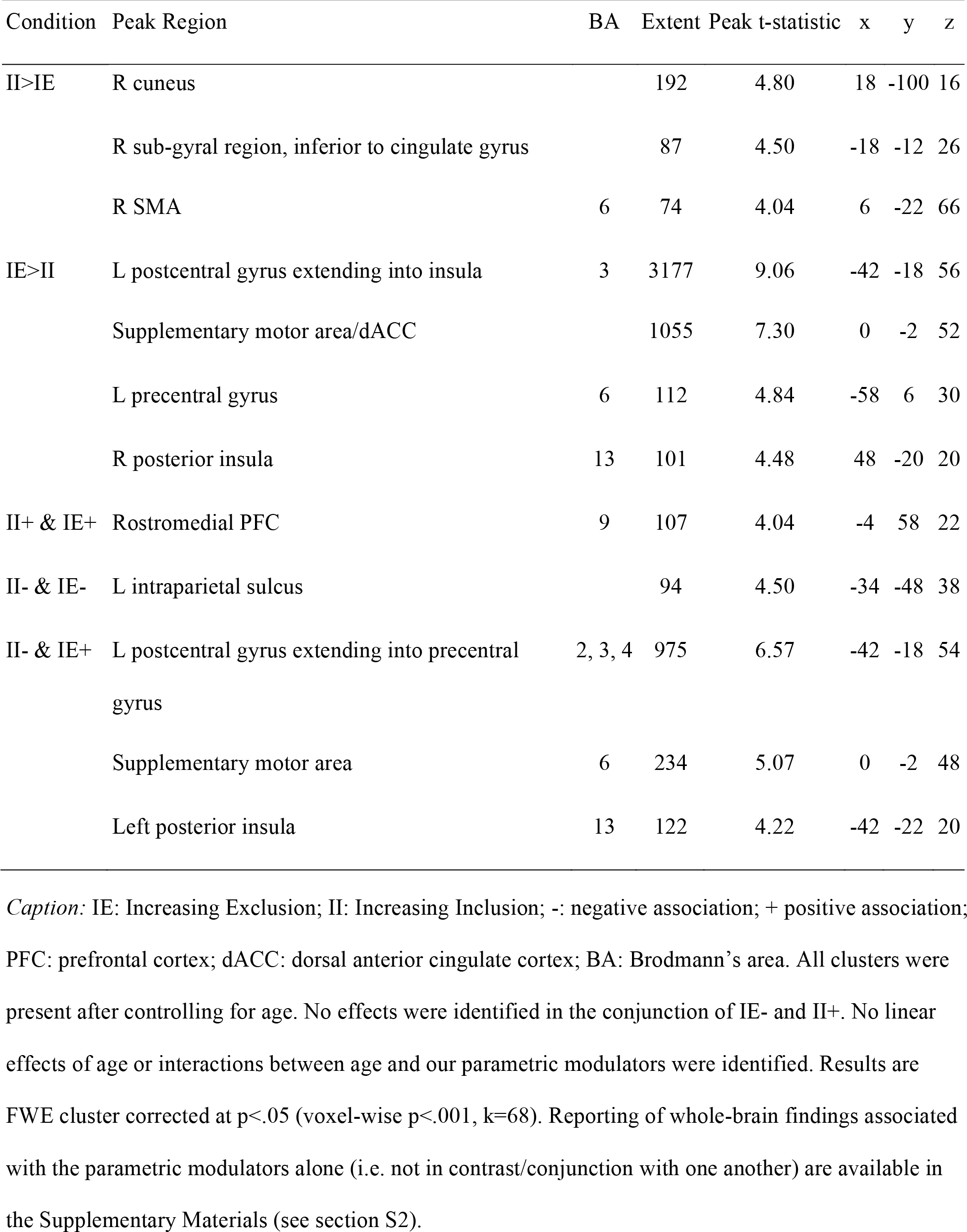
Whole-brain results with parametric modulators

| Condition | Peak Region | BA | Extent | Peak t-statistic | x | y | z |
| --- | --- | --- | --- | --- | --- | --- | --- |
| II>IE | R cuneus |  | 192 | 4.80 | 18 | -100 | 16 |
|  | R sub-gyral region, inferior to cingulate gyrus |  | 87 | 4.50 | -18 | -12 | 26 |
|  | R SMA | 6 | 74 | 4.04 | 6 | -22 | 66 |
| IE>II | L postcentral gyrus extending into insula | 3 | 3177 | 9.06 | -42 | -18 | 56 |
|  | Supplementary motor area/dACC |  | 1055 | 7.30 | 0 | -2 | 52 |
|  | L precentral gyrus | 6 | 112 | 4.84 | -58 | 6 | 30 |
|  | R posterior insula | 13 | 101 | 4.48 | 48 | -20 | 20 |
| II+ & IE+ | Rostromedial PFC | 9 | 107 | 4.04 | -4 | 58 | 22 |
| II- & IE- | L intraparietal sulcus |  | 94 | 4.50 | -34 | -48 | 38 |
| II- & IE+ | L postcentral gyrus extending into precentral gyrus | 2, 3, 4 | 975 | 6.57 | -42 | -18 | 54 |
|  | Supplementary motor area | 6 | 234 | 5.07 | 0 | -2 | 48 |
|  | Left posterior insula | 13 | 122 | 4.22 | -42 | -22 | 20 |

*Caption:* IE: Increasing Exclusion; II: Increasing Inclusion; -: negative association; + positive association; PFC: prefrontal cortex; dACC: dorsal anterior cingulate cortex; BA: Brodmann's area. All clusters were present after controlling for age. No effects were identified in the conjunction of IE- and II+. No linear effects of age or interactions between age and our parametric modulators were identified. Results are FWE cluster corrected at $p < .05$ (voxel-wise $p < .001$ , $k=68$ ). Reporting of whole-brain findings associated with the parametric modulators alone (i.e. not in contrast/conjunction with one another) are available in the Supplementary Materials (see section S2).

#### 3.2.3 Whole-brain responses in the repeated measures 2×2 ANOVA

See Table 4 for comparisons of context congruence, and see section S3 in the Supplementary Materials for detailed whole-brain results from this 2×2 ANOVA. We further explore short-term expectancy violations during Cyberball by examining the interaction between context and event type, which compares when events are congruent (i.e. IncEvent-IncContext and ExcEvent-ExcContext) versus incongruent (i.e. IncEvent-ExcContext and ExcEvent-ExcContext). The contrast of Context Incongruent > Context Congruent events identified the bilateral putamen, bilateral inferior parietal lobule (extending into right posterior superior temporal sulcus and posterior insula), bilateral inferior frontal gyrus, left AI, left precentral gyrus, left precuneus, left posterior superior temporal sulcus, left middle frontal gyrus, right superior temporal gyrus, supplementary motor area, cerebellum, thalamus, and dACC (see Figure 5). Comparing Context Congruent > Context Incongruent events yielded no significant clusters.

**Figure 5.**
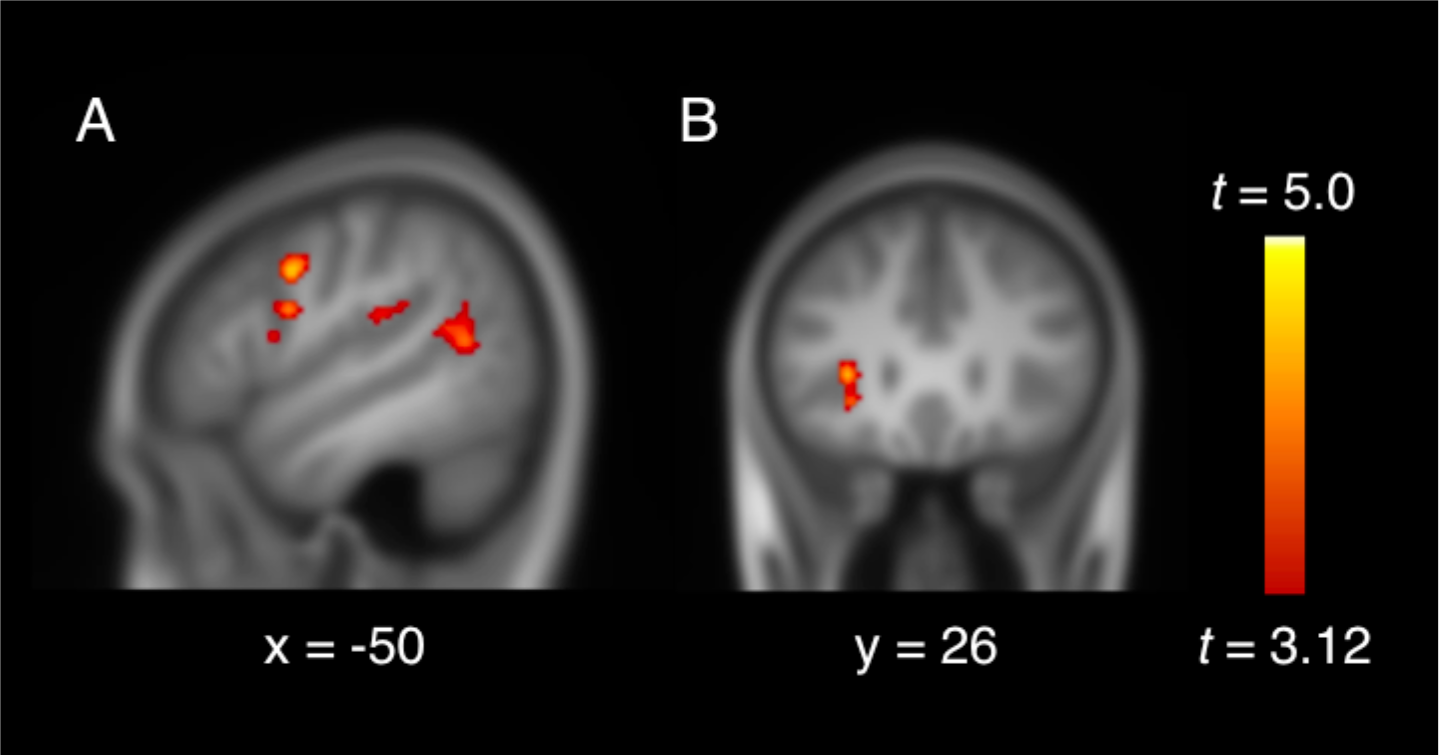
Context Incongruent > Context Congruent results of the event by context interaction. (A) Sagittal view displays the left premotor cortex, posterior insula, and posterior superior temporal sulcus. (B) Coronal view displays the left AI cluster. Results are FWE cluster corrected at p<.05 (voxel-wise p<.001, k=68).

**Table 4.**
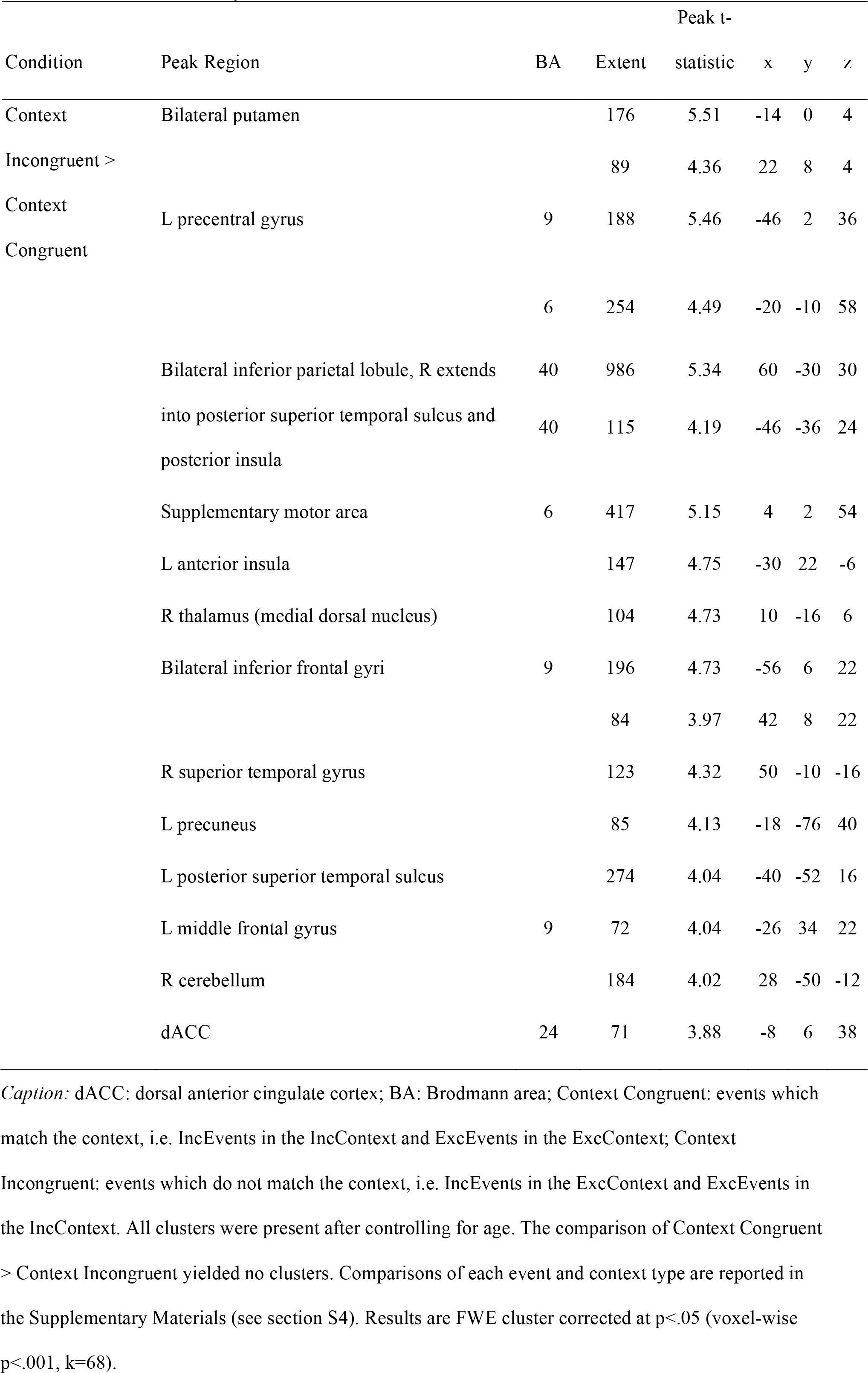
Whole-brain results with 2×2 factorial ANOVA

Age effects and age interactions in the repeated measures 2×2 ANOVA did not reveal clusters in the ACC or insula, and these analyses are reported in the Supplementary Materials (see section S4). Controlling for age did not substantively alter findings in the repeated measures 2×2 ANOVA.

## 4. Discussion

This study characterizes neural scaling to cumulative exclusionary and inclusionary events during Cyberball, focusing on evaluating affective distress models as compared to alternate accounts of dACC and AI functioning. Interestingly, our results support diverse roles for the dACC and insula during Cyberball. Specifically, we found that (a) dACC responses to exclusion are correlated with affective distress, (b) the dACC and posterior insula exhibit signal increases with exclusion as well as *decreases* with over-inclusion, and (c) the dACC and AI respond to events that violate short-term expectancies established by the context of each run. Additionally, we used conjunction analyses to identify regions that respond to both types of social interactions, notably including the left ventrolateral PFC and rostromedial PFC. Finally, we identified a trending negative association between age and responses to cumulative exclusion in the VS that is consistent with age effects in the literature.

### 4.1 Evaluating an affective distress model of dACC and insula functioning

We evaluated the hypothesis that dACC and AI responses to social exclusion during Cyberball reflect affective distress. To fully support an affective distress model, these regions should exhibit greater signal scaling with Increasing Exclusion (cumulative exclusion events in the exclusion context) than Increasing Inclusion (cumulative inclusion events in the inclusion context), and this difference should be attributable to signal increases with Increasing Exclusion only. Whole-brain analyses revealed that the caudal dACC exhibited significantly greater signal in Increasing Exclusion than Increasing Inclusion (ROI analyses were consistent with this finding, but were not significant after correction for multiple comparisons). Exploratory analyses found that affective distress was positively associated with signal responses to Increasing Exclusion in a functionally-defined caudal dACC ROI. However, greater signal in the dACC was driven not only by significant increases in signal with Increasing Exclusion, but also by non-significant but notable (in effect size) *decreases* in signal with Increasing Inclusion. Similarly, a cluster in the left *posterior* insula exhibited significant signal increases with Increasing Exclusion and significant decreases with Increasing Inclusion. These findings intriguingly suggest that the dACC and posterior insula are both involved in processing *over*-inclusion. A prior study implicated both the dACC and AI in social acceptance in addition to rejection (Dalgleish et al., 2017), although these regions responded similarly to both conditions, rather than in opposing directions. Nevertheless, our findings are not fully consistent with an affective distress model, as this model does not predict signal decreases with over-inclusion.

However, one important consideration is that there may be individual differences in affective distress related to being in the scanner environment. For adolescents experiencing such distress, signal decreases with Increasing Inclusion may reflect an attenuation of distress with inclusion over time (cumulative inclusion as “less negative affect”). Future research might clarify this possibility by measuring affective distress at the beginning and end of the inclusion run, but doing in a within-subjects design may draw attention to the manipulation and impact its ecological validity.

### 4.2 Evaluating alternate models of the dACC and insula functioning

The most prominent alternate model of dACC and AI functioning emphasizes the role of these regions in processing expectancy violations (Bush, Luu, & Posner, 2000). We first examined support for this model using parametric modulators. As both exclusion and over-inclusion might similarly violate fair play expectations, an expectancy violation model implies that signal would scale similarly to both types of events. This pattern was not observed in the dACC or AI. However, we also examined neural responses associated with violations of short-term expectancies developed through repeated events of one type establishing a context *within* a run (contrast of Context Incongruent > Context Congruent events). This contrast identified the dACC, left AI, and bilateral posterior insula as processing short-term expectancy violations. These are different regions than those associated with expectancy violations across longer time-scales (i.e. linear change in neural responses to consecutive events using parametric modulators). Interestingly, the dACC and posterior insula clusters associated with context incongruence partially overlapped with those that exhibited greater signal in Increasing Exclusion than Increasing Inclusion. As it seems unlikely that Context Incongruent events elicit distress across both contexts, the involvement of these regions is not necessarily associated with rejection-related processing in social rejection paradigms. Thus we caution against reverse inference of dACC and insula involvement as distress-related, unless otherwise corroborated. Considered together, our two analysis approaches (parametric modulators and unmodulated analysis of congruency between events and context) implicate overlapping regions of the dACC and posterior insula in different processes within the same paradigm.

Other interpretations remain plausible (e.g., participant response demands also scale in opposite directions for cumulative exclusion and inclusion, with cumulative exclusion requiring fewer participant button responses), and our paradigm is unable to disentangle these competing explanations. Future studies with more detailed measures of participants’ expectations of inclusion and degree of affective distress might adjudicate these theories, although it is difficult to do so without conspicuously drawing attention to the manipulation and impacting the psychological experience and believability of the task.

### 4.3 Neural regions underlying both exclusion and over-inclusion

The rostromedial and left ventrolateral PFC exhibited signal increases to *both* Increasing Exclusion and Increasing Inclusion. Prior meta-analyses have consistently implicated the anterior PFC in social exclusion (Cacioppo et al., 2013; Vijayakumar et al., 2017), with recent work indicating that this region processes negative affect (Kragel et al., 2018). Our whole-brain results suggest that this region processes unexpected and/or affectively charged social interaction more generally. Its involvement may reflect self-oriented processing, as the anterior PFC is implicated in perspective-taking (Ames et al., 2008), prospection (Addis et al., 2007) and self-reflection (Modinos et al., 2009). The left ventrolateral PFC ROI encompasses Brodmann area 47 and is distinct from areas of the ventrolateral PFC associated with the regulation of negative affect (Masten et al., 2009; Wager et al., 2008) and rule violation (Bolling et al., 2011b). Instead, this region is primarily associated with language and semantic processing (Petrides, 2016; Ardila, Bernal, & Rosselli, 2017), and has also been implicated in processing others’ intentions (Brunet, Sarfati, Hardy-Baylé, & Decety, 2000), social norm violations (Berthoz, Armony, Blair, & Dolan, 2002) and social punishments (Spitzer, Fischbacher, Herrnberger, Grön, Fehr, 2007). Future studies might explore whether anterior and left ventrolateral PFC indicate the extent to which cumulative exclusionary or inclusionary interactions are self-relevant and/or socially salient.

### 4.4 Age effects

We identified a trending negative association between age and VS responses to Increasing Exclusion. The VS ROI was independently identified for being more reliably recruited during social exclusion in developmental (ages 7-18) than emerging adult samples, and it is notable that a consistent negative effect was identified in our cross-sectional sample of 11 to 17 year olds. Further developmental research would be beneficial to corroborate this effect, which suggests that changes in VS response with age may not be limited to reward processing (Silverman, Jedd, & Luciana, 2015). This region is known for processing both rewarding (Knutson et al., 2000; Sescousse et al., 2013) and aversive stimuli (Jensen et al., 2003; Levita et al., 2009), and increases in VS responses to social stimuli have been associated with reduced susceptibility to peer influences (Pfeifer et al., 2011). We did not find further associations between age and neural responses to either Increasing Exclusion or Increasing Inclusion, despite prior research identifying age-related changes in medial and ventrolateral PFC, among other regions, during social exclusion as compared to fair play (see Bolling et al., 2011a). However, these studies examine different age ranges than ours, and use less stringent thresholding methods for whole-brain analyses.

### 4.5 Strengths and limitations

The current study includes noteworthy methodological strengths. Parametric modulators model exclusion and over-inclusion-related neural changes in gameplay without arbitrary definitions of when these conditions begin and end. In contrast, the literature widely employs block and event-related designs that model social experience assuming binary, static, and independent periods or occurrences. A parametric approach may better capture changes in neural signal when gameplay is more fluid, reflecting naturalistic social interactions that often involve greater ambiguity than typical laboratory tasks. An additional strength is the use of independent ROIs identified via meta-analysis for being reliably recruited in Cyberball paradigms (that use block and event-related designs). These ROIs exhibited signal increases with Increasing Exclusion, providing evidence in favor of the validity of this modeling approach.

Our study also faced several limitations. First, it did not account for possible order effects, as over-inclusion always preceded exclusion. We did not counterbalance because exclusion paradigms can induce distress among adolescents (Masten et al., 2009; Peake, Dishion, Stormshak, Moore, & Pfeifer, 2013; Sebastian et al., 2010), and we sought to minimize the potential carryover of any such negative affect into the subsequent round of Cyberball. We also prioritized maintaining a consistent affective experience overall, because experiencing over-inclusion after a period of exclusion reflects an affective experience that is distinct from the reverse. Relatedly, inclusion runs occurring before and after exclusion runs have been found to elicit different neural responses in Cyberball (White et al., 2013). Some differences between conditions might also be attributed to consistently viewing participant introduction videos and YLG game play in between the social inclusion and exclusion runs. Watching one another’s videos and gameplay prior to exclusion provided a plausible basis for (computer) players to shift toward a negative evaluation of the participant, and was found to be a critical component for the believability of the manipulation during pilot testing.

Another limitation is that the intensity of over-inclusion and exclusion was imbalanced, with a greater average number of cumulative events in Increasing Exclusion (5.97 cumulative exclusion throws in the exclusion context) than in Increasing Inclusion (2.73 cumulative inclusion throws in the inclusion context). Relatedly, there were few incongruent throws in each context, and a particularly low number of inclusion throws in the exclusion context. We thus have relatively less precision in our estimates of some conditions, weakening certain inferences due to varying ability to detect effects. Furthermore, it is unlikely that the true modulation of neural signal driven by social interactions is linear, particularly as interactions become prolonged (i.e. as the number of events becomes large). Our results provide a valuable estimate of linear signal scaling with a low yet affectively meaningful (based on NTS responses) number of events, but do not elucidate the true shape of the trajectory—with an average of less than three consecutive inclusion throws, this was not feasible. Future studies might also require participants’ button press response even when the ball is not thrown to them in order to minimize the influence of asymmetrical motor demands; we modeled participants’ button press as a regressor (as in Bolling et al., 2011a; Bolling et al., 2011b; Bolling, Pelphrey, & Wyk, 2016), but this may not have fully accounted for preparatory motor responses.

Finally, participants completed the Need Threat Scale in reference to the Cyberball game in general (run unspecified), weakening the specificity of associations between this measure and neural responses to exclusion. However, the exclusion run took place at least ten minutes after the inclusion run, was the most recent run when participants were completing the scale, and participants report average levels of need-threat comparable to those in previous studies of exclusion, suggesting that the exclusion run was the reference point for most participants.

### 4.6 Conclusions

We used parametric modulators to examine the specificity of neural responses to social exclusion as compared to over-inclusion during Cyberball. We found BOLD signal increases with cumulative exclusion in the left and right AI (in sensitivity analyses), left inferior frontal gyrus, left posterior cingulate cortex, VS, and across the extent of the ACC. While dACC signal was associated with affective distress, areas within the dACC and insula scaled negatively with cumulative inclusion events, and also responded to violations of short-term expectancies established by the context of each run. As such, our findings caution against interpreting involvement of the dACC and AI as necessarily reflecting aspects of affective distress in social rejection paradigms. The left ventrolateral PFC and rostromedial PFC exhibited similar signal increases in both Increasing Exclusion and Increasing Inclusion, suggesting that these regions play a role in processing events across conditions. Finally, a trending negative age association with cumulative exclusion events in the VS may reflect changes in emotional reactivity and/or regulation across adolescence.

## Open practices statement

None of the analyses were pre-registered. Statistical maps are available on NeuroVault (https://neurovault.org/collections/3794). Preprocessing scripts used for this analysis are available on GitHub at https://github.com/dsnlab/TDS_scripts/tree/cheng_cyb_main/fMRI/ppc/spm/tds2 (SPM scripts) and https://github.com/dsnlab/TDS_scripts/tree/cheng_cyb_main/fMRI/ppc/shell/schedule_spm_jobs/tds2 (shell scripts). High motion volumes were identified using an in-house automated script that is publicly available (Cosme et al., 2018). We refer interested readers to the most recent version (https://github.com/dsnlab/auto-motion), and the branch used in this analysis is available at https://github.com/dsnlab/TDS_scripts/tree/cheng_cyb_main/fMRI/fx/motion/auto-motion.

## Conflict of Interest

The authors declare no competing financial interests.

## Acknowledgements

The authors wish to express gratitude to Rebecca Calcott for consultation on the analysis and to Danielle Cosme for scripts to process motion and to access brain parcellation maps. This work was supported by the grants P50 DA035763 (PIs: Chamberlain and Fisher) and R01 MH107418 (PI: Pfeifer). Author TWC was supported by National Center for Advancing Translational Sciences of the National Institutes of Health under award number TL1TR002371. The content is solely the responsibility of the authors and does not necessarily represent the official views of the National Institutes of Health.

