## Supplementary Materials for "Feeling left out or just surprised? Neural correlates of social exclusion and over-inclusion in adolescence"

### S1. Sensitivity analyses

#### S1.1. Methods

We conducted two sensitivity analyses in order to evaluate the impact of potential confounds on the effects reported in the primary manuscript. In the first sub-sample, we removed the eight subjects that self-reported that they had a psychiatric diagnosis, were taking psychiatric medication, or both. In the second sub-sample, we removed the eight subjects that answered "No" to the question "Did you think the peers could actually see you playing [the driving game]?", as these participants may not have believed the deception ("subsample without disbelief").

For these subjects, we re-ran comparisons across the set of thirteen ROIs. Additionally, we re-ran the parametric modulators and examined results for comparability. Our strategy sought to confirm the presence of clusters from the original analysis in the sensitivity analyses for main comparisons of interest (contrasts between Increasing Inclusion and Increasing Exclusion as well as the conjunction of these parametric modulators). First, we examined all results at the same cluster-based threshold ($p<.001$, $k\geq68$). If any clusters from the original analysis were not present in the sensitivity analysis, we examined the results at a second threshold with a lower voxel-wise threshold and higher cluster-extent threshold ($p<.005$, $k\geq153$; see the section 2.3.2 of the manuscript for how these thresholds were obtained).

#### S1.2 Results from subsample without psychiatric diagnosis or medication

There are few substantive differences in the ROI results when comparing our original findings and the subsample without psychiatric diagnoses or medications (see Tables A and B). These are discussed in the manuscript (section 3.2.1).

Supplementary Table A

*Revised Table 1, no subjects with psychiatric diagnoses and/or medication (N = 61)*

| ROI | Increasing Exclusion M (SE) | $H_0$: Increasing Exclusion = 0 *t* (*p*) | Increasing Inclusion M (SE) | $H_0$: Increasing Inclusion = 0 *t* (*p*) | $H_0$: Increasing Exclusion = Increasing Inclusion *t* (*p*) |
|---|---|---|---|---|---|
| 1 | .06 (.02) | 3.46 (9.87e-4)$^{***s}$ | -.10 (.06) | -1.84 (.071) | 2.89 (5.42e-3)$^{**}$ |
| 2 | .06 (.02) | 3.44 (1.06e-3)$^{**s}$ | -.03 (.07) | -.49 (.629) | 1.41 (.165) |
| 3 | .09 (.02) | 3.88 (2.66e-4)$^{***s}$ | .03 (.09) | 0.29 (.772) | .722 (.473) |
| 4 | .04 (.02) | 1.82 (.074) | .09 (.07) | 1.21 (.233) | -.69 (.49) |
| 5 | .11 (.02) | 4.68 (1.69e-5)$^{***s}$ | .13 (.09) | 1.89 (6.34e-2) | -.59 (.558) |
| 6 | .07 (.02) | 3.41 (1.15e-3)$^{**s}$ | -.02 (.06) | -.37 (.715) | 0.70 (.488) |
| 7 | .07 (.02) | 3.65 (5.58e-4)$^{***s}$ | .14 (.08) | 1.84 (7.08e-2) | -.70 (.485) |
| 8 | .14 (.03) | 4.85 (9.19e-6)$^{***s}$ | .22 (.11) | 2.08 (4.20e-2) | -.83 (.409) |

*Caption*: M=Mean beta, SE= Standard error; $H_0$= null hypothesis; **p<.01, ***p<.001, $^s$: significant at FDR-adjusted *q*=0.05.

Supplementary Table B

*Revised Table 2, no subjects with psychiatric diagnoses and/or medication (N = 61)*

| ROI | Increasing Exclusion M (SE) | $H_0$: Increasing Exclusion = 0 *t* (*p*) | Increasing Inclusion M (SE) | $H_0$: Increasing Inclusion = 0 *t* (*p*) | $H_0$: Increasing Exclusion = Increasing Inclusion *t* (*p*) |
|---|---|---|---|---|---|
| Left inferior frontal gyrus | .10 (.02) | 4.9 (7.69e-6)$^{***s}$ | .18 (.07) | 2.42 (.019) | -1.1 (.276) |
| Left posterior cingulate cortex | .11 (.03) | 4.32 (6.04e-5)$^{***s}$ | .01 (.10) | .12 (.904) | .95 (.344) |
| Ventral striatum | .09 (.03) | 3.46 (1.01e-3)$^{**s}$ | .35 (.11) | 3.35 (1.41e-3)$^{**s}$ | -2.48 (.016) |
| Left anterior insula | .06 (.02) | 3.74 (4.14e-4)$^{***s}$ | .03 (.05) | .62 (.540) | .51 (.613) |
| Right anterior insula | .06 (.03) | 3.71 (4.60e-4)$^{***s}$ | .02 (.04) | .56 (.578) | .82 (.416) |

*Caption*: M=mean beta, SE= standard error; $H_0$= null hypothesis; **p<.01, ***p<.001, $^s$: significant at FDR-adjusted *q*=0.05.

See Table C for a summary of whole-brain findings from the subsample that excluded subjects with psychiatric diagnoses and/or medication. Two clusters presented in the manuscript were not present at either the initial (p<.001, k>68) or secondary (p<.005, k>153) thresholds: 1) for the comparison of Increasing Exclusion > Increasing Inclusion, the cluster in the left precentral gyrus was no longer present at either threshold, and 2) for the conjunction of signal negatively associated with both Increasing Inclusion and Increasing Exclusion, the cluster in the left intraparietal sulcus was absent. This suggests that contributions from participants with psychiatric diagnoses and/or medication drove the significance of these findings in the original analyses.

Several novel clusters were also identified in these analyses. Notably, some parts of the rostromedial PFC were identified in Increasing Inclusion > Increasing Exclusion. This cluster exhibits partial overlap with a cluster identified from the conjunction of Increasing Inclusion and Increasing Exclusion, suggesting that signal is significantly greater than baseline for *both* conditions, but is even higher for cumulative inclusion events in this subsample for some subregions. Altered modulation of signal in this region in response to social exclusion has been found with respect to one type of psychopathology (schizophrenia; see Gradin et al., 2012). Although we not sufficiently powered to investigate such effects, this finding raises the possibility that signal in this region is altered in other forms of psychopathology as well.

The left insula posterior cluster identified from the conjunction of signal negatively associated with Increasing Inclusion and positively associated with Increasing Exclusion exhibited a slight difference in peak coordinate and was reduced in spatial extent compared to the corresponding cluster in the original analysis.

Supplementary Table C

*Revised Table 3, no subjects with psychiatric diagnoses and/or medication (N = 61)*

| Condition | Peak Region | BA | Extent | Peak t-statistic | x | y | z |
|---|---|---|---|---|---|---|---|
| II>IE | dmPFC | 8 | 95 | 4.43 | 2 | 44 | 50 |
| | R sub-gyral region, inferior to cingulate gyrus | | 140 | 4.36 | -18 | -12 | 26 |
| | R cuneus | 18 | 157 | 4.22 | 20 | -98 | 18 |
| | Caudate extending into ACC [p,**] | | 184 | 4.40 | -16 | 30 | 8 |
| | L cerebellum[p,**] | | 176 | 4.29 | -50 | -64 | -30 |
| | SMA[p,*] | 6 | 253 | 4.15 | 12 | -20 | 72 |
| | Rostromedial PFC [p,**] | 9 | 181 | 3.63 | 12 | 62 | 16 |
| IE>II | L postcentral gyrus extending into insula | 3 | 2088 | 7.80 | -42 | -18 | 56 |
| | SMA/dACC | | 950 | 7.10 | -2 | -2 | 52 |
| | L insula[**] | 13 | 348 | 5.57 | -48 | -18 | 20 |
| | | 13 | 88 | 4.78 | -34 | -2 | 14 |
| | R posterior insula | 13 | 88 | 4.25 | 48 | -20 | 20 |
| | R caudate extending into putamen[**] | | 135 | 4.67 | 20 | 2 | 14 |
| II+ & IE+ | L middle temporal gyrus[**] | 21 | 78 | 4.03 | -58 | -12 | -12 |
| | Rostromedial PFC | 9 | 80 | 4.04 | -4 | 58 | 22 |
| II- & IE+ | L postcentral gyrus extending into precentral gyrus | 2, 3, 4 | 864 | 5.96 | -40 | -18 | 56 |
| | Supplementary motor area | 6 | 294 | 5.30 | 0 | -2 | 50 |
| | Left anterior insula[*] | 13 | 76 | 4.22 | -46 | -20 | 20 |

IE: Increasing Exclusion; II: Increasing Inclusion; -: negative association; +: positive association; [p]: Cluster identified at a lower voxel-wise/higher cluster extent threshold; [*]Cluster is slightly dissimilar—peak coordinates are greater than +/- 2 mm in each of the x, y, or z directions and/or appear different upon visual inspection; ** Novel cluster not present in original analyses

**S1.3 Results from subsample without disbelief.**

There are a few substantive differences in our ROI results when comparing our original findings and the subsample without those that did not believe the manipulation (see Tables D and E). These are discussed in the main manuscript (section 3.2.1).

Supplementary Table D

*Revised Table 1, no subjects indicated that they did not believe the manipulation (N = 61)*

| ROI | Increasing Exclusion M (SE) | $H_0$: Increasing Exclusion = 0 *t* (*p*) | Increasing Inclusion M (SE) | $H_0$: Increasing Inclusion = 0 *t* (*p*) | $H_0$: Increasing Exclusion = Increasing Inclusion *t* (*p*) |
|---|---|---|---|---|---|
| 1 | .07 (.02) | 3.33 (1.49e-3)$^{**s}$ | -.10 (.05) | -1.90 (6.26e-2) | 2.91 (5.05e-3)$^{**}$ |
| 2 | .06 (.02) | 3.51 (8.68e-4)$^{***s}$ | -.04 (.07) | -.60 (.550) | 1.56 (.123) |
| 3 | .09 (.02) | 3.88 (2.65e-4)$^{***s}$ | .01 (.09) | 0.11 (.911) | .88 (.380) |
| 4 | .04 (.02) | 2.07 (4.29e-2) | .06 (.08) | .78 (.437) | -.23 (.822) |
| 5 | .12 (.02) | 4.76 (1.25e-5)$^{***s}$ | .15 (.09) | 1.71 (9.28e-2) | -.27 (.710) |
| 6 | .07 (.02) | 3.26 (1.85e-3)$^{**s}$ | 2.23e-3 (.07) | .03 (.974) | 0.96 (.340) |
| 7 | .07 (.03) | 2.59 (1.19e-2) | .10 (.08) | 1.19 (.237) | -.31 (.756) |
| 8 | .13 (.03) | 4.54 (2.77e-5)$^{***s}$ | .21 (.11) | 1.89 (6.31e-2) | -.72 (.474) |

*Caption*: M=mean beta, SE= standard error; $H_0$= null hypothesis; **p<.01, ***p<.001, $^{s}$: significant at FDR-adjusted *q*=0.05.

Supplementary Table E

*Revised Table 2, no subjects indicated that they did not believe the manipulation (N = 61)*

| ROI | Increasing Exclusion M (SE) | $H_0$: Increasing Exclusion = 0 *t* (*p*) | Increasing Inclusion M (SE) | $H_0$: Increasing Inclusion = 0 *t* (*p*) | $H_0$: Increasing Exclusion = Increasing Inclusion *t* (*p*) |
|---|---|---|---|---|---|
| Left inferior frontal gyrus | .09 (.02) | 4.37 (5.08e-5)$^{***s}$ | .18 (.07) | 2.47 (.016) | -1.29 (.200) |
| Left posterior cingulate cortex | .13 (.03) | 4.39 (4.59e-5)$^{***s}$ | .02 (.10) | .20 (.842) | 1.00 (.322) |
| Ventral striatum | .10 (.03) | 3.43 (1.08e-3)$^{**s}$ | .28 (.11) | 2.42 (.018) | -1.57 (.123) |
| Left anterior insula | .06 (.02) | 2.92 (4.91e-3)$^{**s}$ | .03 (.06) | .45 (.654) | .63 (.531) |
| Right anterior insula | .06 (.02) | 3.13 (2.70e-3)$^{**s}$ | .02 (.04) | .51 (.613) | .89 (.377) |

*Caption*: M=mean beta, SE= standard error; $H_0$= null hypothesis; **p<.01, ***p<.001, [s]: significant at FDR-adjusted *q*=0.05.

See Table F for a summary of whole-brain findings from the subsample that excluded subjects that did not believe the manipulation. We found that all clusters from the original comparisons of Increasing Inclusion and Increasing Exclusion, as well as their conjunctions, were present at either the initial (p<.001, k>68) or secondary (p<.005, k>153) thresholds. Two clusters differed somewhat in their peak coordinates (the right cuneus in Increasing Inclusion > Increasing Exclusion and the left precentral gyrus from the reverse contrast), but upon visual inspection were very similar in the anatomical regions they included.

Supplementary Table F.

*Revised Table 3, no subjects indicated that they did not believe the manipulation (N = 61)*

| Condition | Peak Region | BA | Extent | Peak t-statistic | x | y | z |
|---|---|---|---|---|---|---|---|
| II>IE | R cuneus[*] | | 165 | 4.60 | 12 | -100 | 14 |
| | R sub-gyral region, inferior to cingulate gyrus[p] | | 295 | 3.86 | -18 | -12 | 26 |
| | R SMA[p] | 6 | 167 | 4.04 | 6 | -22 | 66 |
| | L caudate extending into ACC[p,**] | | 165 | 4.13 | -16 | 30 | 8 |
| IE>II | L postcentral gyrus extending into insula | 3 | 3330 | 8.82 | -40 | -18 | 54 |
| | SMA/dACC | | 1141 | 7.18 | 0 | -2 | 52 |
| | L precentral gyrus[*] | 6 | 115 | 4.84 | -52 | 4 | 32 |
| | R precentral gyrus[**] | 6 | 73 | 4.27 | 32 | -10 | 66 |
| | R posterior insula | 13 | 76 | 4.33 | 48 | -20 | 20 |
| | L cerebellum[**] | | 84 | 4.04 | -20 | -82 | -12 |
| II+ & IE+ | Rostromedial PFC[p] | 9 | 254 | 4.04 | -4 | 56 | 22 |
| II- & IE- | L intraparietal sulcus | | 271 | 4.14 | -34 | -48 | 38 |
| | L inferior parietal[**] | 40/7 | 222 | 3.70 | -40 | 52 | 56 |
| II- & IE+ | L postcentral gyrus extending into precentral gyrus | 2, 3, 4 | 1014 | 6.48 | -40 | -18 | 54 |
| | Supplementary motor area | 6 | 272 | 5.01 | 0 | -2 | 48 |
| | Left anterior insula | 13 | 226 | 4.98 | -42 | -22 | 20 |

IE: Increasing Exclusion; II: Increasing Inclusion; -: negative association; +: positive association [p]: Cluster from modified p<.005; [*]Cluster differs somewhat from a cluster by the same name in the original analysis on the basis of a difference in peak coordinate (greater than +/- 2 mm in each of the x, y, or z directions); ** Novel cluster not present in original analyses

**S2. Whole-brain findings related to the parametric modulators**

Clusters associated with the Increasing Exclusion and Increasing Inclusion parametric modulators are displayed below in Table G. All of the clusters presented in this table are present after controlling for age.

Supplementary Table G.

*Whole-brain results with parametric modulators*

| Condition | Peak Region | BA | Extent | Peak t-statistic | x | y | z |
|---|---|---|---|---|---|---|---|
| II+ | L sub-gyral region extending into mid-cingulate gyrus | | 162 | 4.67 | -18 | -12 | 26 |
| | R cuneus | | 83 | 4.45 | 18 | -100 | 18 |
| | L middle temporal gyrus | 21 | 105 | 4.31 | -66 | -20 | -12 |
| | Dorsomedial PFC | | 151 | 4.31 | 2 | 44 | 50 |
| | Rostromedial PFC | 9 | 387 | 4.25 | 0 | 60 | 14 |
| | L inferior frontal gyrus | 47 | 84 | 3.82 | -42 | 30 | -14 |
| | L superior frontal gyrus/frontal eye fields | | 68 | 3.86 | -14 | 56 | 34 |
| II- | Bilateral postcentral gyri | 3,7 | 2809 | 7.43 | -42 | -18 | 56 |
| | | 5, 7 | 142 | 3.98 | 32 | -40 | 68 |
| | Medial frontal gyrus, extending into cingulate gyrus | 6 | 542 | 6.21 | -2 | -2 | 52 |
| | L precentral gyrus | 6 | 224 | 5.67 | -58 | 6 | 30 |
| | L insula | 13 | 102 | 4.96 | -36 | -4 | 14 |
| | L supramarginal gyrus | | 96 | 4.79 | -36 | -50 | 38 |
| IE+ | L postcentral gyrus | 3, 1 | 1248 | 6.74 | -42 | -18 | 52 |
| | Paracentral lobule extending into medial frontal and cingulate gyri | 31, 6, 24 | 1290 | 5.96 | -2 | -12 | 46 |
| | Middle temporal gyrus | | 1186 | 5.94 | -50 | -4 | -16 |
| | Rostromedial PFC extending into perigenual ACC | 32 | 1238 | 5.78 | -4 | 58 | -2 |
| | Bilateral insula | | 403 | 5.34 | 36 | 8 | 10 |
| | | 13 | 441 | 5.00 | -38 | -22 | 20 |
| | L anterior middle temporal gyrus | 38, 2 | 92 | 4.97 | -44 | 0 | -42 |
| | R superior temporal gyrus | 42 | 177 | 4.73 | 66 | -24 | 12 |
| | L posterior cingulate | | 344 | 4.70 | -4 | -60 | 12 |

| | | | | | | | |
|---|---|---|---|---|---|---|---|
| | Bilateral inferior frontal gyrus | | 270 | 4.68 | 50 | 26 | 0 |
| | | | 90 | 3.89 | -54 | 24 | 14 |
| | L posterior middle temporal gyrus | | 181 | 4.38 | -68 | -48 | -2 |
| | L angular gyrus | | 114 | 4.28 | -46 | -78 | 30 |
| | R occipitotemporal area/parahippocampal gyrus | 37 | 70 | 5.44 | 32 | -46 | -8 |
| IE- | Precuneus extending into bilateral superior and inferior parietal lobe | | 7420 | 7.78 | 4 | -62 | 58 |
| | Bilateral middle frontal gyrus extending into R dorsolateral PFC | | 3522 | 7.17 | 26 | 2 | 60 |
| | | 6 | 655 | 6.44 | -26 | 0 | 60 |
| | L dorsolateral PFC | 9 | 1250 | 7.13 | -46 | 30 | 36 |
| | Cerebellum | | 1592 | 6.01 | -36 | -64 | -42 |
| | | | 233 | 5.70 | -34 | -42 | -44 |
| | | | 77 | 4.16 | 28 | -38 | -42 |
| | R inferior temporal gyrus | | 74 | 3.90 | 46 | -66 | 0 |

*Caption:* ACC: anterior cingulate cortex, PFC: prefrontal cortex, IE: Increasing Exclusion; II: Increasing Inclusion; -: negative association; +: positive association

**S3. Whole-brain results from the 2x2 factorial ANOVA**

Clusters associated comparisons of event and context in the 2x2 ANOVA are displayed in the table below.

Table H.

*Whole-brain results with 2x2 factorial ANOVA*

| Condition | Peak Region | BA | Extent | Peak t-statistic | x | y | z |
|---|---|---|---|---|---|---|---|
| ExcEvent> IncEvent | L middle occipital gyrus extending into bilateral temporal-occipital cortex | 19 | 11332 | 8.78 | -50 | -72 | 10 |
| | Bilateral inferior parietal lobule | | 2997 | 6.25 | -46 | -38 | 60 |
| | | | 834 | 4.77 | 40 | -40 | 44 |
| | Bilateral middle frontal gyrus | 6 | 4629 | 5.83 | 30 | 20 | 56 |

| | | | | | | | |
|---|---|---|---|---|---|---|---|
| | | 46 | 772 | 5.39 | -48 | 48 | 2 |
| | L orbitofrontal cortex | | 90 | 5.60 | -24 | 32 | -24 |
| | L posterior insula | | 163 | 5.41 | -40 | -12 | 16 |
| | L postcentral gyrus/superior temporal gyrus | 40 | 127 | 5.30 | -60 | -24 | 14 |
| | L postcentral gyrus | 3 | 165 | 5.05 | -42 | -18 | 40 |
| | Temporal pole extending into middle temporal gyrus | 21 | 271 | 5.13 | 40 | 12 | -46 |
| | L parahippocampal gyrus extending into superior temporal gyrus | 20 | 112 | 4.72 | -26 | -4 | -44 |
| | R supramarginal gyrus | 22/40 | 186 | 4.30 | 64 | -44 | 40 |
| | R uncus* | 20 | 68 | 4.19 | 30 | -6 | -42 |
| | R dorsolateral PFC | 9 | 123 | 3.84 | 46 | 26 | 32 |
| IncEvent > ExcEvent | L precentral gyrus | | 1231 | 8.62 | -34 | -18 | 52 |
| | R medial frontal gyrus* | 6 | 395 | 6.03 | 16 | -12 | 66 |
| | R thalamus | | 72 | 3.76 | 10 | -12 | -4 |
| ExcContext >IncContext | R precentral gyrus | | 194 | 4.74 | 54 | -2 | 16 |
| | | 4 | 157 | 4.73 | 52 | -4 | 42 |
| | R inferior parietal lobule | | 236 | 4.16 | 58 | -22 | 24 |
| | L posterior insula | | 103 | 3.83 | -42 | -18 | 20 |
| IncContext> ExcContext | Bilateral inferior parietal lobule | 40 | 545 | 4.99 | -48 | -34 | 46 |
| | | | 269 | 4.82 | 42 | -38 | 40 |
| | Bilateral superior and middle frontal gyri | 6 | 317 | 4.66 | -24 | 0 | 56 |
| | | | 328 | 4.47 | 24 | -2 | 52 |
| | Bilateral precuneus | | 233 | 4.43 | 18 | -72 | 56 |
| | | 7 | 182 | 4.25 | -18 | 68 | 58 |

*Caption:* PFC: prefrontal cortex; IncEvent: inclusion event, i.e. throw to the participant; ExcEvent= exclusion event, i.e. throw to computer player; IncContext: inclusion context, ie., the inclusion run; ExcContext: exclusion context, i.e. the exclusion run; Context Congruent: events which match the context, i.e. IncEvents in the IncContext and ExcEvents in the ExcContext; * Cluster is no longer present when controlling for age. Results are FWE cluster corrected at $p<.05$ (voxel-wise $p<.001$, $k=68$).

**S4. Age-related analyses in the repeated measures 2x2 ANOVA**

Using whole-brain regression, we examined associations with age in both the 2x2 ANOVA and in the model with parametric modulators. We were interested in interactions between the linear age term and the context and throw factors in the 2x2 ANOVA, as well as between the linear age term and the parametric modulators.

A number of age-related findings from the repeated measures 2x2 ANOVA are reported in Supplementary Table I. These findings bear little resemblance to developmental differences identified from the meta-analysis (Vijayakumar et al., 2017), which may be less surprising after taking into account that meta-analysis findings were derived from group comparisons of developmental (child/adolescent) and emerging adult samples, whereas the current analyses examine age-associations within the adolescent period. However, our findings also bear little resemblance to continuous linear age associations identified in work by Bolling and colleagues (2011). There are a myriad of differences between these studies and the present study, including the schedule of throws and modeling strategies, which makes it difficult to explain inconsistent findings. Additionally, there are a number of Cyberball studies in child and adolescent samples that do not report age effects. More widespread reporting of such effects, even when they are not central to the questions of interest, may better help us understand developmental patterns in neural responses to Cyberball.

Supplementary Table I.

*Whole-brain results with age in the repeated measures 2x2 ANOVA*

| Condition | Peak Region | BA | Extent | Peak t-statistic | x | y | z |
|---|---|---|---|---|---|---|---|
| Age+ | Bilateral middle occipital gyri | | 509 | 6.95 | -44 | -82 | 4 |
| | | 18, 37 | 159 | 4.94 | 56 | -70 | -2 |
| | | | 81 | 4.15 | 34 | -94 | 8 |
| | Superior frontal gyrus | 6 | 165 | 4.67 | 4 | 16 | 64 |
| | R cerebellum | | 235 | 4.47 | 26 | -94 | -18 |
| | R transverse temporal gyrus | 42 | 75 | 4.41 | 68 | -14 | 14 |
| | R cerebellum | | 75 | 4.31 | 0 | -78 | -22 |
| | L precentral gyrus | 6 | 156 | 4.29 | -42 | -2 | 54 |
| | Bilateral cuneus | | 133 | 4.26 | 2 | -86 | 30 |
| Age by Throw | L primary motor cortex | | 124 | 4.21 | -32 | -20 | 50 |

+: positive association. No clusters were negatively associated with age, nor were they associated with the age by context interaction. The left primary motor cortex was associated with an interaction between age and throw type. Visual inspection of the parameter estimates from this cluster suggests that this region exhibits greater average signal during exclusion throws (throws to the computer) in older participants.

### S5. Smoothing

We also sought to determine if signal in the functionally defined dACC cluster (4 mm radius sphere around MNI coordinates -8, 6, 38) had been amplified by signal from the nearby supplementary motor cortex due to smoothing. This concern was minimized as we found the same pattern of ROI results in a separate unsmoothed model.
